# CSDC2, an RBP essential to cardiomyocyte commitment during cardiac differentiation in hiPSC

**DOI:** 10.64898/2026.02.12.705579

**Authors:** Rubens Gomes-Júnior, Isabela Tiemy Pereira, José Henrique Rosa da Silva, Giovanna Nazaré Prezia, Lucia Spangenberg, Tamara Fernandez-Calero, Bruno Dallagiovanna

## Abstract

Cardiovascular diseases are the leading cause of death worldwide, accounting for approximately 30% of total mortality. Changes in post-transcriptional regulation have been correlated with the development of cardiopathies. RNA-binding proteins (RBP) are proteins capable of interacting with mRNAs, regulating their stability, localization, and translation. Here, we described CSDC2 as an RBP expressed at the final stages of cardiac differentiation using hPSCs as a model. We showed that the loss of CSDC2 impairs cardiomyocyte differentiation, while the recovery of its expression rescues the differentiation potential of these cells. We characterized the translatome of CSDC2 knockout cells during cardiac differentiation by polysome profiling. In cardiac mesoderm cells, CSDC2 interacts with ribosomal proteins. Furthermore, CSDC2 appears to be able to associate with mRNAs encoding regulators of cardiac progenitor commitment. Altogether, in this study, we describe a new role of CSDC2 in cardiomyocyte commitment using cardiac differentiation of hiPSCs.

## Introduction

RNA-binding proteins (RBPs) are proteins capable of interacting with RNA to form ribonucleoprotein complexes, thereby influencing RNA structure and its interactions with other molecules^1^. RBPs are involved in several steps of post-transcriptional regulation (PTR) of RNA metabolism, acting as regulators of RNA biogenesis, stability, function, transport, and localization^2,3^. Due to their central role in PTR, RBPs have been described as key players in various biological processes, including cell proliferation, cell cycle progression, migration, and differentiation^4–6^. Moreover, dysregulation of RBPs is associated with a wide range of pathological conditions^7^.

Cardiovascular diseases remain the leading cause of death worldwide, according to the latest data from the World Health Organization^8^. Since the discovery of human pluripotent stem cells (hPSCs), these cells have become a valuable tool for studying cardiac development and cardiopathies^9^. Cardiac differentiation of hPSCs has been widely employed to identify molecules involved in normal cardiac development, which may contribute to the establishment of more robust differentiation models or the discovery of molecular dysfunctions potentially associated with cardiac diseases^10^.

Over the years, the importance of RBPs in developmental processes, maintenance, and homeostasis of cardiomyocytes has been recognized^11–14^. A correlation between PTR dysregulation and the development of cardiac pathologies has been observed in studies reporting altered patterns of mRNA polyadenylation and splicing in samples from patients with cardiac hypertrophy and congenital heart disease^15–17^. Several RBPs have been identified as key regulators of heart development. Among them, RBM20 stands out as a well-characterized RBP in the cardiac context. RBM20 has been shown to regulate the splicing of more than 40 RNA targets, and its dysfunction is associated with dilated cardiomyopathy^18,19^. Another example is the Quaking protein (QKI), which regulates essential genes in cardiomyocytes, such as *ACTN2*, *MYO18A*, and *HEY*^20,21^. Another RBP involved in cardiac development is RBM24, which is expressed in the human fetal heart, and loss-of-function assays in zebrafish have demonstrated that its depletion reduces sarcomeric protein levels, resulting in impaired cardiac contractility^22,23^. These examples highlight the critical role of RBPs in heart development and the pathogenesis of cardiac diseases.

Previous data from our group exploring cardiac differentiation using embryonic stem cells revealed several RBP genes being translated^24^. Among them is Cold Shock Domain-Containing C2 (*CSDC2*), also named as PIPPin^25^. CSDC2 protein is composed of two putative RNA-binding domains, PIP1 and PIP2, which flank its central cold shock domain^25^. CSDC2 has been described as participating in the polyadenylation of histone mRNAs during neural development in rats^26,27^. In vivo studies using a rat model of decidualization have shown increased expression of CSDC2 and changes in its subcellular localization during this process; cytoplasmic localization is associated with differentiated tissue, whereas nuclear localization is observed in proliferative regions^28^. CSDC2 has also been linked to certain types of cancer, such as carcinoma and glioblastoma, where reduced expression levels have been reported^29,30^. Conversely, CSDC2 has been proposed as a potential marker for prostate cancer^31^.

Here, we describe the expression profile of CSDC2 during cardiac differentiation using both hESC and hiPSC models. We demonstrate that the lack of expression of CSDC2 impairs hiPSC-derived cardiomyocyte differentiation, and that expression recovery rescues the differentiation capacity. Using proximity labeling and immunoprecipitation techniques, we characterize the interactome of CSDC2 and identify potential cellular pathways regulated by this protein during the differentiation process. Finally, we identified some RNAs that CSDC2 could interact with during cardiac differentiation.

## Methods

### Cell Culture

Human embryonic stem cells, H1 line, were obtained from WiCell Research Institute (Madison, WI, USA) under a Materials Transfer Agreement (No. 18-W0416) with the Carlos Chagas Institute. hiPSC line IPRN13.13 was kindly provided by the Kyba Lab at the University of Minnesota, USA ^32^. Both hESC and hiPSC lines are derived from a male and exhibit a regular karyotype. The cells were cultured on Geltrex™-coated dishes (Gibco) using Stem Flex™ medium (Gibco). Passage was performed using Accutase™ (Gibco) after the culture reached 80% confluence. Cells were maintained at 37°C with a 5% CO2 atmosphere.

### Cardiac Differentiation

The differentiation protocol was adapted from Lian^33^. The cells were seeded at 10^5^ cells/cm^2^ in a Geltrex-coated 24-well plate using Stem Flex medium with 10 μM Y27632 (Med Chem Express). For the next two days, the medium was daily replaced until the culture reached 100% confluence. The cells were induced to mesoderm in 2 mL of RPMI 1640 (Gibco) supplemented with 2% B-27 minus insulin (Gibco) and 12 μM CHIR99021 (TOCRIS). The next day, 24 h after mesoderm induction, the medium was replaced with 2 mL of RPMI supplemented with 2% B-27 minus insulin. On day 3 of differentiation, 1 mL of conditioned medium was mixed with 1 mL of fresh RPMI supplemented with 2% B-27 minus insulin medium and 10 µM XAV939 and added to the well. On day 5, the medium was replaced with 2 mL of fresh RPMI supplemented with 2% B-27 minus insulin, and on days 7, 10, and 13, the medium was replaced with 2 mL of RPMI supplemented with 2% B-27. Spontaneous beating starts at day 11.

### Immunofluorescence

Cells were fixed with 4% paraformaldehyde (PFA) for 15 minutes at room temperature (RT), then washed twice with phosphate-buffered saline (PBS). Subsequently, the cells were permeabilized using PBS containing Triton X-100 (0.3%) for 30 minutes at RT, followed by blocking with PBS containing bovine serum albumin (BSA) (1%) for 1 hour at RT. The primary antibodies: anti-TNNT2 (Abcam - ab45932 – 1:100); anti-CSDC2 (Santa Cruz - sc-376693 – 1:100); anti-OCT4 (Abcam - ab19857 – 1:100); anti-SSEA4 (Abcam - ab16287 – 1:100) were diluted in PBS/BSA, then incubated with the cells for 16 hours at 4°C. Next, the cells were washed three times with PBS, followed by incubation with anti-mouse, rabbit, or goat IgG Alexa Fluor™ 488 or 546 (Invitrogen – A11003, A11008, A11010, A21202, A1117036 - 1:1,000) diluted in PBS/BSA for 1 hour at RT. After another round of washing, the cells were incubated with 4’,6-diamidino-2-phenylindole (DAPI, 300mM) for 20 minutes at RT. Following staining, the plate was stored at 4°C for up to 14 days. The images were acquired using fluorescence microscopy (Leica DMI6000B) and analyzed with LAS AF software.

### CRISPR design and knockout cell line

To knock out the CSDC2 gene in the hiPSC, we used the double nicking strategy with the pSpCas9n(BB)-2A-GFP (PX461) vector constructed with two guide RNAs targeting the start codon region on the CDS. The gRNAs were designed by the UCSC Genome Browser and CHOPCHOP web tool, and the off-targets were evaluated by the Cas-OFFinder web tool. The vectors were transfected using Lipofectamine Stem Transfection Reagent (Invitrogen - STEM00008) with 500 ng of each vector per well in a 24-well plate. The mutants were sorted using a BD FACSAria™ II Flow Cytometer, and the genomic edition was screened by PCR.

### Cytometry

hiPSC and hiPSC-derived cardiomyocytes were dissociated using trypsin-EDTA (0.05%) and fixed using 4% paraformaldehyde and 90% methanol. Fixed cells were incubated with 0.5% PBS/BSA, 0.1% Triton-X, and primary antibodies for TNNT2 (Thermo Scientific™ - MS-295-P0 – 1:100) or CSDC2 (Santa Cruz - sc-376693 – 1:100). After washing, cells were incubated with 0.5% PBS/BSA, 0.1% Triton-X, and anti-mouse IgG Alexa Fluor™ 546 (Invitrogen - A11003 – 1:1,000). Analyses were carried out using a FACSCanto II flow cytometer (BD Biosciences) and FlowJo software (v. 10.7).

### Polysome profiling

Polysome profiling was performed as previously described^34^. Cells at differentiated stages were treated with 0.1 mg/mL cycloheximide for 10 min at 37 °C. After incubation, cells were detached and lysed in polysome lysis buffer (15 mM Tris-HCl, pH 7.4, 15 mM MgCl2, 0.3 M NaCl, 1% Triton X-100, 40 U/μL RNAse-Out, 50 U/μL DNAse). Lysis was performed for 10 min on ice, then centrifuged at 12,000 xg for 10 min at 4 °C. The supernatants were loaded on 10–50% sucrose gradients (BioComp model 108 Gradient Master v.5.3) and centrifuged (SW40 rotor, HIMAC CP80WX HITACHI) at 270,000 × g for 120 min at 4 °C. Sucrose gradient fractions were isolated using the ISCO gradient fractionation system (ISCO Model 160 Gradient Former Foxy Jr. Fraction Collector) coupled to a UV light for RNA detection, which recorded the polysome profiling at 254 nm. Fractions were identified based on the peak pattern observed in the RNA detection, the polysome fractions were pooled and added to an equal volume of TRI Reagent (Merck - T9424).

### RNA isolation and qPCR

Total RNA and polysome RNA were isolated by Direct-zol™ RNA kit (Zymo® Research), following the manufacturer’s instructions. cDNA synthesis was performed using ImProm-II Reverse Transcription System (Promega - A3800), following the manufacturer’s instructions. Quantitative PCR reactions were performed with GoTaq® qPCR Master Mix (Promega). Analyses were carried out in the QuantStudio ™ 5 Real-Time PCR System.

### RNA-seq

Polysome RNA-seq was performed with the hiPSC wild-type and hiPSC CSDC2 KO on day 5 and day 10 from the cardiac differentiation protocol. For the RNA-pulldown sequencing, the RNA was isolated from the pulldown using the CSDC2 recombinant or empty resin. Library preparation was performed with TruSeq Stranded mRNA (Illumina – 20020594). Quality assessment and trimming of reads were done with FastQC and Trim Galore (v.0.4.0). Reads were mapped to hg38/GRCh38 with HISAT2 (v.2.1.0) and counted with HTSeq (v.0.11.1). For the translatome RNAseq, significantly differentially expressed genes were identified with DESeq2 (v.1.24.0) with an adjusted p-value cutoff of 0.05 and a log2FoldChange cutoff of |2|. For pulldown RNA-seq, the RNA sequencing raw read counts were normalized using CPM-TMM. The normalized values for each condition were summarized using the median expression across biological replicates to reduce noise caused by replicate variability and outliers’ influence. After that, we computed the top quantile threshold (top 5%) of median log2FC values to identify the strongest pulldown-enriched candidates. Peak calling was conducted using Piranha (http://smithlab.usc.edu), and binding motif analysis was performed using HOMER software and Multiple Em for Motif Elicitation (MEME - Version 5.5.9). For confirmation of motifs, we used the FIMO tool from MEME suite.

### Proximity labeling

For the proximity labeling, we cloned the CSDC2 sequence into a TurboID backbone vector (Addgene #226995)^35^. The hiPSC with the inducible-expression system of CSDC2-TurboID and GFP control (Addgene #226995) was submitted to the cardiac differentiation protocol. The TurboID expression was induced for 4 hours with 1 μg/mL doxycycline. The biotinylated proteins were purified using High-Capacity Magne® Streptavidin Beads (Promega - V7820). Streptavidin-bead slurry was washed twice with binding/washing buffer (25 mM Tris-HCl, 150 mM NaCl, pH 7.5). Then, the total protein extract from the cells was incubated with the beads for 16 hours at 4°C. After binding, the beads were washed three times with binding/washing buffer. For elution, the beads were resuspended in sample buffer (300 mM Tris-HCl, 0.3 M SDS, 0.1M DTT, 14 mM β-mercaptoethanol, 6% glycerol, and 0.05% bromophenol blue) and heated at 98°C for 15 minutes. Eluted proteins were analyzed by liquid chromatography–mass spectrometry (LC-MS).

### Immunoprecipitation

Immunoprecipitations were performed using Dynabeads™ Protein G (Invitrogen - 10003D). hiPSC with an inducible-expression system for CSDC2 were submitted to the cardiac differentiation protocol. 16 hours before the immunoprecipitation, the cells were induced with 1 μg/mL doxycycline. On the following day, cells were lysed in NP-40 lysis buffer (50 mM Tris-HCl, 150 mM NaCl, 1% NP-40) containing protease inhibitor (Roche - 11836170001) for 30 minutes at 4 °C. Lysates were then centrifuged at 13,000 xg for 15 minutes, and the supernatants were collected. Protein G beads were prepared with either anti-CSDC2 antibody (Santa Cruz – sc376693) or mouse IgG isotype control (Merck - PP54) according to the manufacturer’s instructions. Protein extracts were incubated with the beads for 2 hours at room temperature. Beads were then washed three times with PBS. Bound proteins were eluted in sample buffer (described above) and heated at 98 °C for 10 minutes. Eluted proteins were analyzed by liquid chromatography–mass spectrometry (LC-MS).

### LC-MS and proteomic analysis

Affinity-purified peptides were analyzed using the Ultimate 3000 RSLCnano system coupled to the Orbitrap Exploris 120 mass spectrometer (Thermo Fisher Scientific). Label-free quantification (LFQ) data were analyzed using Perseus software^36^. For statistical analysis, LFQ values were log2-transformed, and missing values were removed. These values were subjected to a Student’s t-test comparing CSDC2 and control in the day 5 and day 10 samples. Volcano plots were generated using −log of p-value and Log2ΔLFQ. The cutoff for positive interaction peptides was set at p-value<0.05.

### Pull-down

Before the RNA-pulldown assay, the CSDC2 recombinant was expressed in *E. coli* (BL21-pLysS) for 5 hours at 30°C. The protein was purified by affinity chromatography using ÄKTA pure (Cytiva). The RNA was obtained from hiPSC differentiated to mesoderm cardiac using the same cardiac differentiation protocol. For the pull-down, 50 µg of CSDC2 recombinant was immobilized using Ni-NTA (Qiagen - 30210). The resin was blocked by incubating with PBS with 1%BSA and RNAseOUT for 1 hour. Next, 10 µg of the RNA isolated was incubated with immobilized CSDC2 for 1 hour at room temperature, followed by three washes with PBS. The RNA capture by CSDC2 was eluted in 500 µL of TRI Reagent (Merck - T9424), followed by RNA isolation.

### Statistical analysis

Graphed data are expressed as mean ± SD. Statistical analysis was performed with Prism 8.0 software (GraphPad Software). For comparisons between two mean values, the student’s unpaired t-test was performed. A p-value lower than 0.05 was considered statistically significant. Frequency of beating areas was counted by collaborators blinded to the experimental conditions.

## Results

### CSDC2 expression is increased in the final stages of cardiac differentiation

In previous work, our group described the gene expression pattern in a human embryonic bodies’ differentiation model (Fig. 1A), and characterized the translatome of this process using the polysome-profiling^24^. Here, we checked this dataset to look for promising candidates of RBP as regulators of cardiac differentiation. We found the RBP CSDC2 as expressed after mesoderm commitment to cardiomyocyte differentiation (Fig. 1B). We confirmed this expression profile in different human pluripotent stem cells (hPSCs) and using a monolayer cardiac differentiation protocol (Fig 1C). When both human embryonic stem cells (hESC) and human induced pluripotent stem cells (hiPSC) were used, we detected an increased CSDC2 expression in the final stages of cardiac differentiation: cardiac progenitor cells (day 10) and early cardiomyocytes (day 15) (Fig. 1D). Next, we confirmed the presence of CSDC2 in cardiac progenitors and cardiomyocytes at the protein level, and observed that the protein is localized mainly in the cytoplasm (Fig. 1E). For further experiments, we use the monolayer protocol due to its robustness and less complexity.

**Fig. 1.**
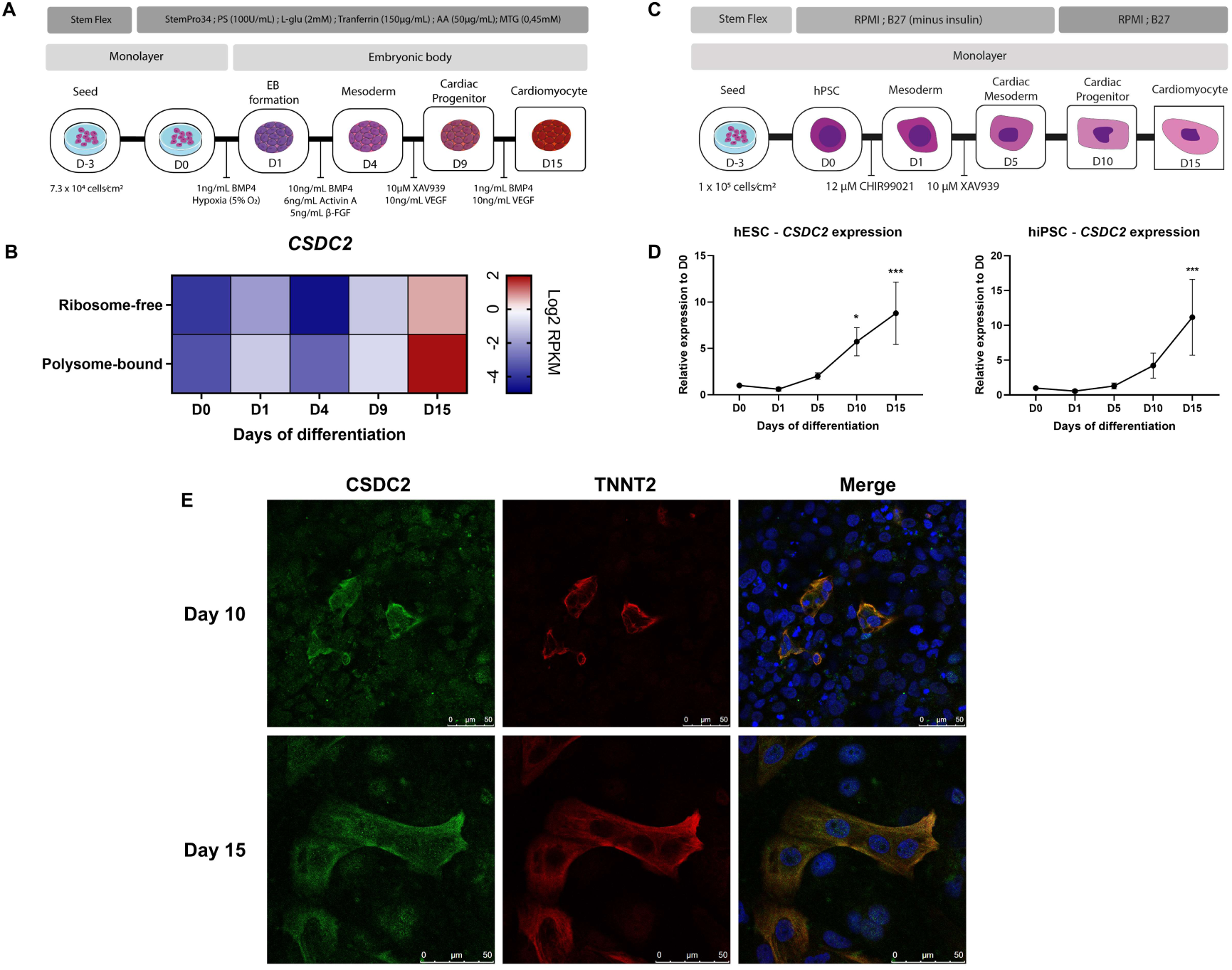
Characterization of CSDC2 expression during cardiomyocyte differentiation of hPSCs. **A.** Schematic representation of cardiac differentiation using an embryoid body formation protocol. **B.** RNA-seq data of CSDC2 expression at different time points in ribosome-free and polysome-bound samples during cardiac differentiation through embryoid body formation. **C.** Schematic representation of cardiac differentiation using a monolayer differentiation protocol. **D.** CSDC2 expression during cardiac differentiation using the monolayer protocol in hESCs (left panel) and hiPSCs (right panel), assessed by qPCR. Mean with SD; one-way ANOVA followed by Dunnett’s test. Each column compared with D0. *p≤0.05; *** p≤0.001. **E.** Immunofluorescence analysis of CSDC2 at the cardiac progenitor (day 10) and cardiomyocyte (day 15) stages during cardiac differentiation using the monolayer protocol. CSDC2 is shown in green, TNNT2 in red, and nuclei in blue. Scale bars, lower right.

### CSDC2 is necessary for cardiomyocyte differentiation in hiPSCs

To investigate the relevance of CSDC2 in cardiac differentiation, we generated a CSDC2 knockout hiPSC line (hiPSC KO CSDC2) using the double-nicking strategy described by Ran et al. (2013)^37^ , and targeted indels in the CSDC2 gene near the start codon (Fig. 2A). We screened several clones and identified one with an eight-nucleotide deletion by Sanger sequencing (Fig. 2B). This deletion resulted in a frameshift that altered the amino acid sequence after position 13 and generated a stop codon at position 71 (Fig. S1A). The deletion was confirmed by PCR using a primer designed to anneal directly to the deletion region; amplification occurred only in wild-type (WT) cells, confirming the homozygous mutation in the KO line (Fig. S1B). We differentiated the KO hiPSCs into cardiac progenitors (day 10) and did not detect CSDC2 protein in these cells compared to WT, confirming the knockout at the protein level (Fig. 2C). Additionally, we characterized the KO cells and observed normal colony formation, no chromosomal alterations, and sustained expression of pluripotency markers at both mRNA and protein levels (Fig. S1C–F).

**Fig. 2.**
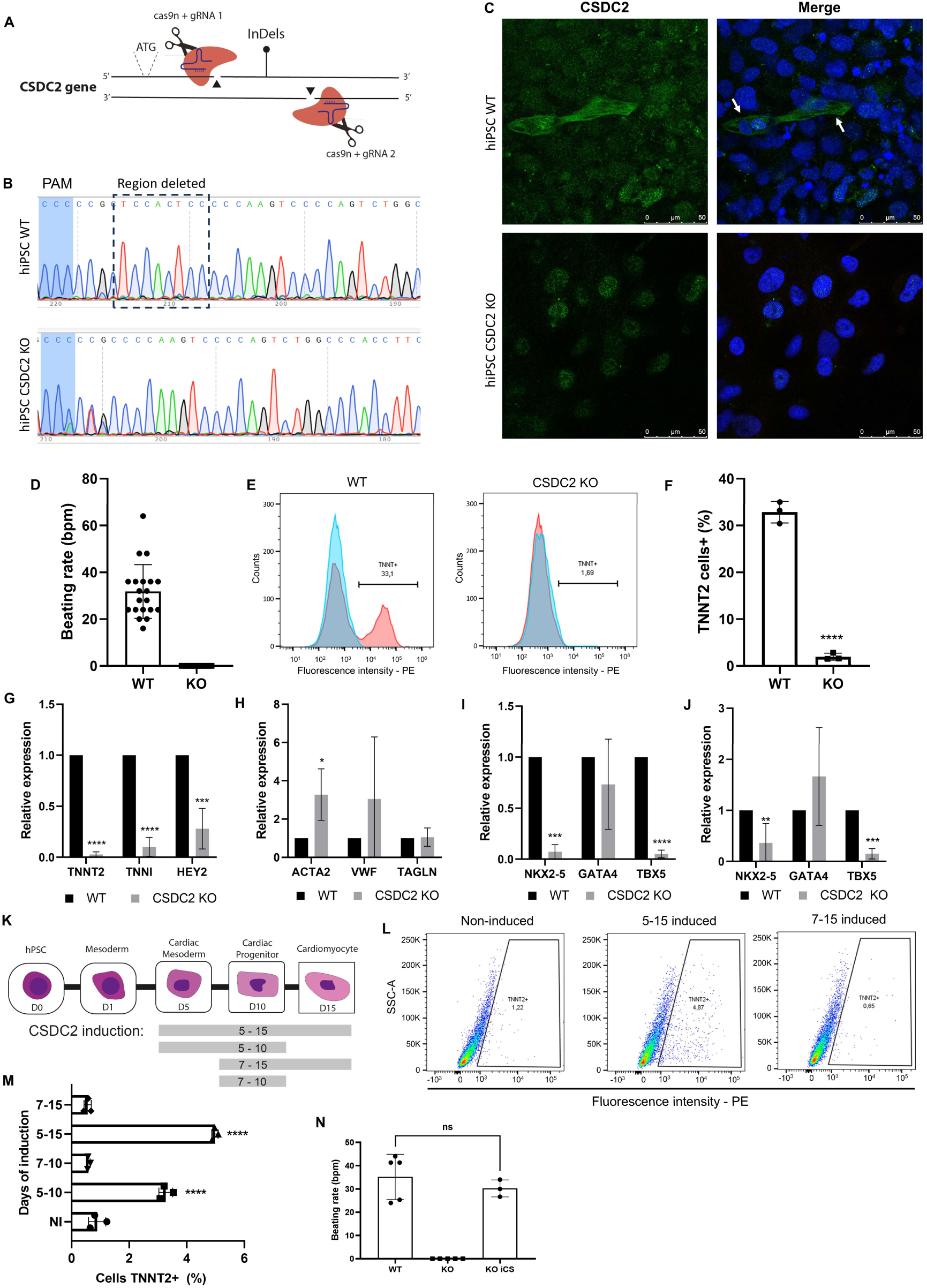
Knockout of CSDC2 impairs cardiomyocyte differentiation in hiPSCs. **A.** Schematic representation of the CRISPR knockout strategy. Arrowheads indicate the cut region. The start codon is shown. **B.** Sanger sequencing of hiPSC wild-type (WT) (upper panel) and CSDC2 knockout (KO) (lower panel). The deleted region is highlighted in WT by a dashed square. **C.** Immunofluorescence of CSDC2 at the cardiomyocyte stage (day 15) during cardiac differentiation using the monolayer protocol. CSDC2 is shown in green, and nuclei in blue. hiPSC WT (upper panel) and CSDC2 KO (lower panel). **D.** Beating rates of cardiomyocytes derived from hiPSC WT and CSDC2 KO. **E.** Flow cytometry analysis of TNNT2-positive cells after cardiac differentiation using hiPSC WT (left panel) and hiPSC CSDC2 KO (right panel). **F.** Statistical analysis of cardiac differentiation efficiency in WT and CSDC2 KO cells. The y-axis represents the percentage of TNNT2-positive cells. **G.** Relative expression of cardiomyocyte markers in hiPSC WT and CSDC2 KO at day 15 of the cardiac differentiation protocol, assessed by qPCR. For qPCR statistical, Mean with SD, Student’s T test. *p≤0.05; **p≤0.01; ***p≤0.001; ****p≤0.0001. The legend is on the right side. **H.** Relative expression of other cell-type markers in hiPSC WT and CSDC2 KO at day 15 of the cardiac differentiation protocol, assessed by qPCR. **I.** Relative expression of master regulators of the cardiomyogenic process in hiPSC WT and CSDC2 KO at day 15 of the cardiac differentiation protocol, assessed by qPCR. **J.** Relative expression of master regulators of the cardiomyogenic process in hiPSC WT and CSDC2 KO at day 10 of the cardiac differentiation protocol, assessed by qPCR. **K.** Schematic representation of the timing of CSDC2-induced expression in hiPSC KO iCSDC2 during cardiac differentiation. **L.** Flow cytometry analysis of TNNT2-positive cells after cardiac differentiation, comparing no induction, 5–15 days of induction, and 7–15 days of induction. **M.** Statistical analysis of TNNT2-positive cells after cardiac differentiation at different times of CSDC2 induction for the recovery phenotype. **N.** Beating rate of hiPSC WT, CSDC2 KO, and KO iCSDC2 (recovery phenotype, induction for 5-15 days) at day 15 of the cardiac differentiation protocol.

Both KO and WT cells were subjected to our cardiac differentiation protocol. KO cells did not exhibit spontaneous beating, in contrast to WT cells (Fig. 2D). To explore whether this phenotype was due to alterations in electrophysiology or differentiation efficiency, we compared the proportion of TNNT2-positive cells in both populations. TNNT2-positive cells were absent in KO cultures (Fig. 2E–F). Consistently, other cardiomyocyte markers, such as TNNI and HEY2, were also reduced in KO cells (Fig. 2G), suggesting that these cells failed to properly differentiate into cardiomyocytes. We then investigated if CSDC2 loss affected other cell types that typically emerge at day 15 in our protocol, such as endothelial cells, smooth muscle cells, or fibroblasts. The expression of ACTA2, VWF, and TAGLN was maintained or even increased in KO cells (Fig. 2H), indicating that CSDC2 knockout selectively impairs cardiomyocyte differentiation. To gain mechanistic insight, we assessed the expression of three core regulators of cardiomyocyte differentiation (Fig. 2I). NKX2-5 and TBX5 expression were reduced in KO cells, whereas GATA4 remained unchanged. The same pattern was observed at the progenitor stage (day 10), when these genes begin to be expressed (Fig. 2J). In contrast, markers of mesoderm and cardiac mesoderm remained at similar or slightly higher levels in KO cells (Fig. S1G-H). Together, these results show that CSDC2 loss reduces the expression of core cardiomyocyte regulators (NKX2-5 and TBX5) from the progenitor stage onward, leading to downregulation of cardiomyocyte-specific genes without affecting other cardiac-derived cell types or mesoderm commitment.

### Recovery of CSDC2 expression in KO cells rescues cardiomyocyte differentiation capacity

To confirm that the alterations observed in cardiac progenitors were exclusively due to CSDC2 loss, we constructed an integrative cassette for inducible expression of CSDC2 (Fig. S2A). This system was introduced into KO CSDC2 cells, and we evaluated its functionality and the proportion of cells responding to induction (Fig. S2B-C). CSDC2 expression was detected only in the presence of the inducer (doxycycline) with approximately 60% of cells expressing the protein distributed in the cytoplasm. The insertion of the cassette did not alter the expression of pluripotency markers in these cells (Fig. S2D), which are subsequently referred here as hiPSC KO iCSDC2.

To assess if the recovery of CSDC2 expression would rescue the cardiomyocyte differentiation and determine a specific time window required to do so, we tested four induction periods during cardiac differentiation: days 5–10, 5–15, 7–10, and 7–15 (Fig. 2K). Interestingly, induction from days 5–10 and 5–15 partially restored cardiomyocyte differentiation capacity, whereas induction from days 7–10 or 7–15 did not (Fig. 2L–M). CSDC2 starts to be expressed from day 5 of differentiation in WT cells (Fig. 1D), which could explain the importance of this time window. We then differentiated hiPSC KO iCSDC2 and induced CSDC2 expression on days 5-15 to evaluate their functionality. Remarkably, KO iCSDC2 cells rescued their spontaneous beating capacity, with a beat rate comparable to WT (Fig. 2N). These results show that the re-introduction of CSDC2 expression in KO cells was able to rescue both cardiomyocyte differentiation and spontaneous beating, and it is specifically required from day 5 of the differentiation onward.

### Translatome of hiPSC-derived cardiac progenitors is affected by CSDC2 KO

Since we confirmed the relevance of CSDC2 during cardiomyocyte differentiation, we next looked for alterations in gene expression in knockout cells. For this, we performed RNA sequencing of the translatome at two time points of differentiation, days 5 and 10. The polysome profiling technique was used to isolate RNAs under translation; at this point, no differences were observed in the polysome profile patterns when comparing hiPSC KO CSDC2 to WT (Fig. S3A). On the other hand, PCA showed clear clustering of each group, suggesting differences between WT and CSDC2 KO data in the gene expression profile (Fig. S3B). Gene expression analysis revealed unique patterns at both time points when comparing WT and KO CSDC2 (Fig. 3A). We defined differentially expressed genes (DEGs) as those with log2 fold change (FC) ≥ |2| and FDR ≤ 0.05. In this case, a positive FC indicated enrichment in WT cells, while a negative FC indicated enrichment in KO cells (Fig. 3B). At the cardiac mesoderm stage (day 5), we observed 708 DEGs enriched in WT and 575 enriched in KO; at the cardiac progenitor stage (day 10), we found 942 DEGs enriched in WT and 660 enriched in KO. To describe the cell profile of these samples, we ranked the DEGs found in cardiac progenitors and performed gene set enrichment analysis (GSEA). This showed a positive score (genes enriched in WT) for cardiac cell development and cardiomyocyte differentiation, and a negative score (genes enriched in KO) for processes related to smooth muscle and epithelium (Fig. 3C). Gene Ontology (GO) analysis revealed dysregulation in KO cells starting from the cardiac mesoderm stage. DEGs enriched in WT cells were associated with terms such as “Mesoderm development” and “Regulation of heart contraction,” whereas DEGs enriched in KO cells were linked to terms such as “Renal sodium excretion” and “Neuron differentiation” (Fig. 3D). Furthermore, human tissues GO analysis at the cardiac progenitor stage indicated that WT samples were enriched for gene sets related to heart and cardiovascular system, while KO cells resembled plasma cells and bone marrow cells (Fig. 3E). Together, these data demonstrate that CSDC2 knockout alters the translatome during cardiac differentiation in hiPSCs. Moreover, KO cells lose the expression of many genes normally present in cardiac mesoderm and cardiac progenitors, a phenomenon that may explain their inability to differentiate into cardiomyocytes.

**Fig. 3.**
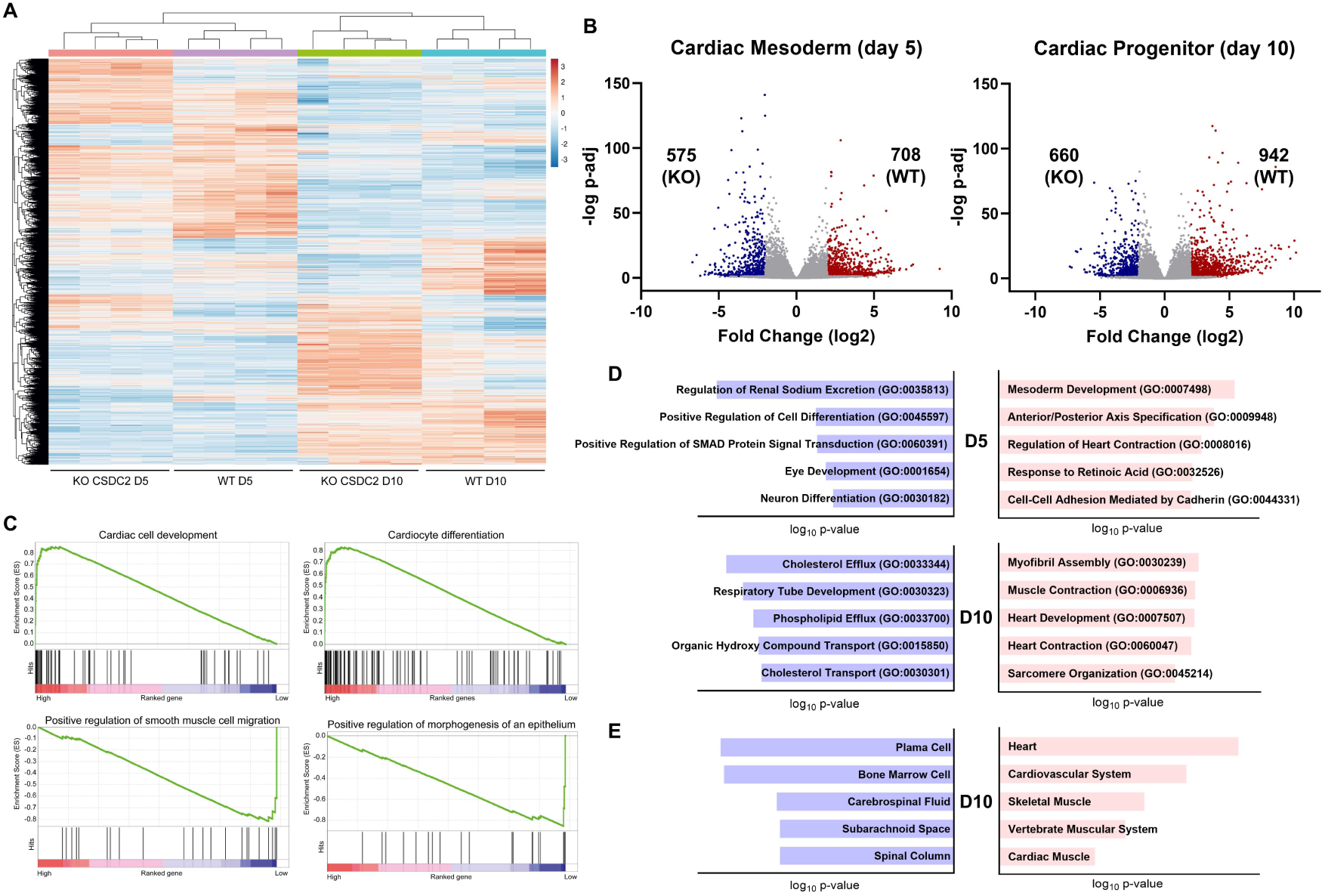
Translatome analysis of CSDC2-knockout cells at cardiac mesoderm and progenitor stages. **A.** Heatmap of the translatome profile from hiPSC WT and CSDC2 KO at days 5 and 10 of the cardiac differentiation protocol. Values are presented as normalized read counts. The color key is displayed to the right of the chart. **B.** Volcano plots for cardiac mesoderm (day 5) and cardiac progenitor (day 10) showing data from hiPSC WT and CSDC2 KO cells. Blue dots represent genes enriched in CSDC2 KO cells, while red dots indicate genes enriched in WT cells compared to pluripotent hESCs. The number of genes is indicated above the dots. **C.** Gene set enrichment analysis (GSEA) of RNA-seq data from the cardiac progenitor stage (day 10). The four enrichment plots were arbitrarily selected from the top twenty enriched gene sets to represent two positive and two negative enrichment scores. **D.** Gene Ontology analysis for biological processes in cardiac mesoderm (day 5) and cardiac progenitor (day 10). Blue charts represent genes enriched in CSDC2 KO cells, and red charts represent genes enriched in WT cells. **E.** Gene Ontology analysis for human tissues in the cardiac progenitor stage (day 10). Blue charts represent genes enriched in CSDC2 KO cells, and red charts represent genes enriched in WT cells.

### CSDC2 interacts with ribosomal proteins in the cardiac mesodermal stage

To describe the mechanism of CSDC2 in the cardiac context, we identified its interaction partners during our cardiac differentiation protocol. First, we employed the proximity labeling technique using the inducible expression system previously constructed and validated^35^. In this system, CSDC2 was engineered as a fusion with TurboID, a biotin ligase (Fig. S4A). We verified the system functionality in the presence of doxycycline and determined the minimum induction time required to detect CSDC2 expression (Fig. S4B–C). After these standardizations, hiPSC iCSDC2–TurboID cells were subjected to our differentiation protocol until day 5 or day 10. At each stage, CSDC2–TurboID expression was induced for 4 hours, and the biotinylated proteins were identified by mass spectrometry (MS) (Fig. 4A). The same system with GFP–TurboID served as a control for promiscuous biotinylation.

**Fig. 4.**
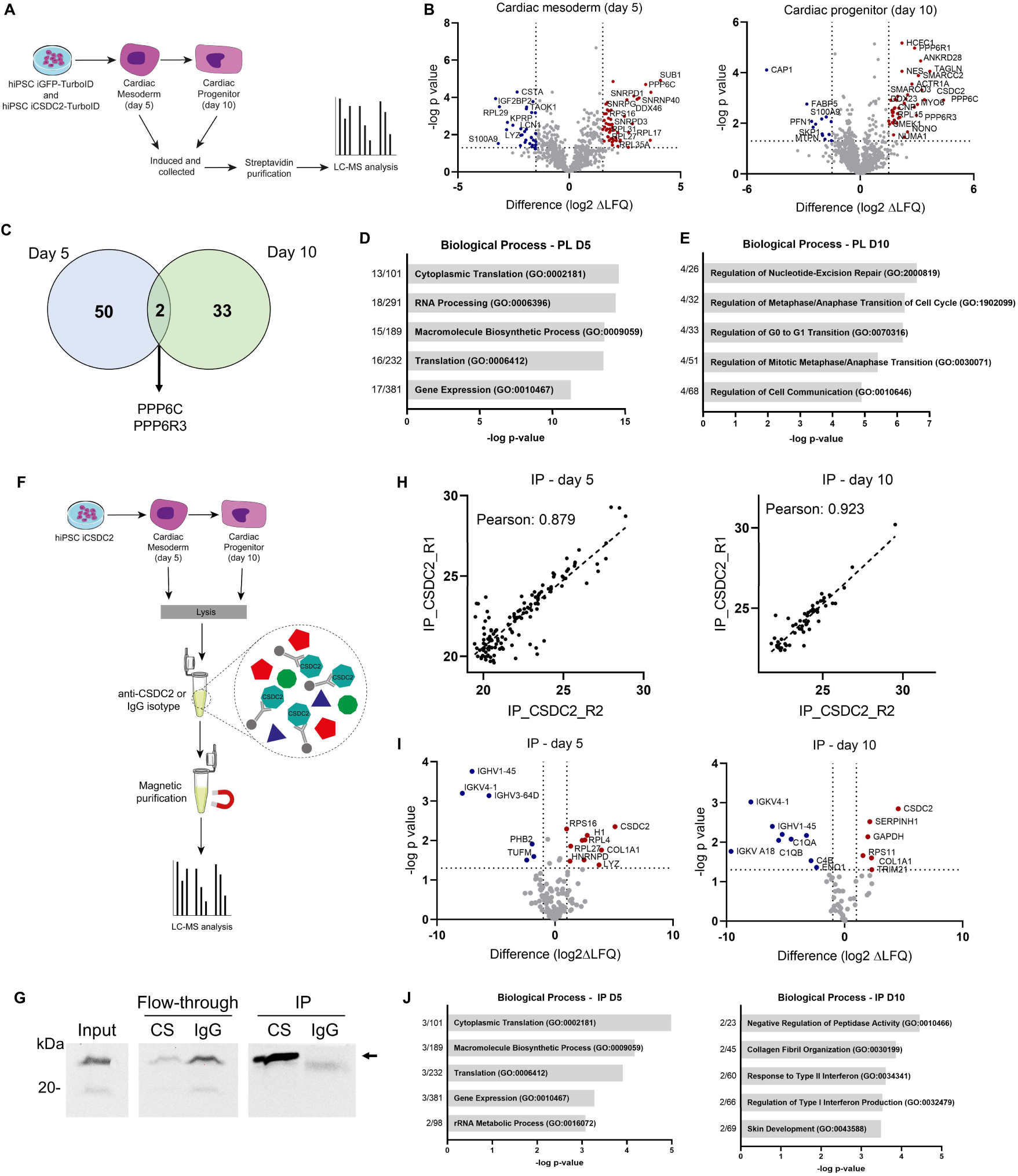
Description of the CSDC2 interactome from proximity labeling and immunoprecipitation. **A.** Experimental design for CSDC2 proximity labeling. **B.** Volcano plots for cardiac mesoderm (day 5 -left) and cardiac progenitor (day 10 - right) showing data from hiPSC iCSDC2-TurboID and iGFP-TurboID. Blue dots represent proteins enriched in the GFP control, while red dots indicate proteins enriched in CSDC2 samples. Y-axis threshold: p-value ≤ 0.05 (−log p-value ≥ 1.3); x-axis threshold: log2 LFQ difference ≥ 1.5. Gene names are shown next to their representative dots. **C.** Venn diagram of proteins identified by proximity labeling of CSDC2 at the cardiac mesoderm (day 5) and cardiac progenitor (day 10) stages. Proteins common to both datasets are shown below. **D–E.** Gene Ontology analysis of biological processes for proteins identified as interacting with CSDC2 in cardiac mesoderm (day 5 - D) and cardiac progenitor (day 10 - E). **F.** Experimental design for CSDC2 immunoprecipitation. **G.** Western blot analysis of the immunoprecipitation assay. CS, CSDC2; IgG, mouse isotype IgG; IP, immunoprecipitation. The arrow indicates the representative CSDC2 band. **H.** Pearson correlation coefficient analysis of CSDC2 immunoprecipitation in cardiac mesoderm (day 5 -left) and cardiac progenitor (day 10 -right). **I.** Volcano plots of immunoprecipitation data for CSDC2 and IgG control. Blue dots represent proteins enriched in the IgG control, while red dots indicate proteins enriched in CSDC2 samples. Y-axis threshold: p-value ≤ 0.05 (−log p-value ≥ 1.3); x-axis threshold: log2 LFQ difference ≥ 1.5. Datasets from cardiac mesoderm (day 5 - left) and cardiac progenitor (day 10 - right) are shown. **J.** Gene Ontology analysis of biological processes for proteins identified in CSDC2 immunoprecipitation at the cardiac mesoderm (day 5 - left) and cardiac progenitor (day 10 - right).

Principal component analysis (PCA) showed that induced iCSDC2–TurboID, non-induced (NI), and GFP–TurboID samples formed distinct clusters at both differentiation stages (Fig. S4D-E). Initially, we compared peptides identified in induced vs. non-induced CSDC2–TurboID samples. This analysis revealed unspecific biotinylation of chaperone complex components, probably due to overexpression of the exogenous protein (Fig. S4F-G), and these peptides were excluded from further analyses. We then compared CSDC2–TurboID with GFP–TurboID to identify genuine CSDC2 interactors at the cardiac mesoderm and cardiac progenitor stages (Fig. 4B). We found a distinct set of proteins at each stage: in cardiac mesoderm, several translation-related proteins such as DDX46, RPS16, RPL17, RPL27, and RPL31; and in cardiac progenitors, a protein complex involved in cell cycle regulation—PPP6C, PPP6R1, PPP4R1, PPP6R3—along with a few translation-related proteins such as DDX23 and RPL15. Notably, only PPP6C and PPP6R3 were present in both datasets (Fig. 4C), suggesting interactions that may occur at both differentiation stages. Gene Ontology (GO) analysis revealed distinct enrichment profiles: cardiac mesoderm interactors were associated with PTR terms such as “Cytoplasmic translation,” “RNA processing,” and “Gene expression,” whereas cardiac progenitor interactors were enriched for cell cycle-related terms such as “Regulation of metaphase/anaphase transition” and “Regulation of G0 to G1 transition” (Fig. 4D-E).

To validate these interaction patterns, we performed immunoprecipitation (IP) as a complementary approach. Due to the low endogenous expression of CSDC2, we used our inducible expression system to overexpress CSDC2 in WT cells, followed by its immunoprecipitation and its partners identification by MS (Fig. 4F). We confirmed the functionality of the system; the proportion of cells responding to induction and validated the IP protocol by western blot (Fig. S5A-B; Fig 4G). We also tested whether CSDC2 overexpression enhanced differentiation efficiency in WT cells, but no significant increase was observed (Fig. S5C). In PCA analysis, samples from the IP of CSDC2 and the Isotype control formed distinct clusters for both cardiac mesoderm (D5) and cardiac progenitor (D10) (Fig. S5 D-E). Pearson correlation coefficients for the IP experiments exceeded 0.85, indicating high sample similarity and robustness of the technique (Fig. 4H). In the cardiac mesoderm stage, IP recovered proteins related to gene expression, such as RPS16, RPL4, RPL27, and HNRNPD, while in cardiac progenitors, only RPS11 was identified (Fig. 4I). GO analysis of IP-identified partners recapitulated the proximity labeling results at the cardiac mesoderm stage, with enrichment for PTR-related terms such as “Cytoplasmic translation,” “Gene expression,” and “Translation” (Fig. 4J). These results strongly suggest the role of CSDC2 as a RBP during the cardiac differentiation. On the other hand, terms involving matrix organization appeared in the analysis of progenitor stage, different from proximity labeling results. Together, these results highlight the dynamic, stage-dependent interactome of CSDC2 during cardiac differentiation, suggesting that its RNA-binding activity is more prominent in the cardiac mesoderm stage.

### CSDC2 can regulate cardiac progenitor differentiation by mRNA interactions

To better understand the CSDC2 mechanism, we investigated its potential RNA targets. To achieve this, we performed a pull-down (PD) assay using the recombinant His-CSDC2 to capture RNAs that interact with the protein from cardiac mesoderm samples, followed by RNA sequencing. First, we verified the individuality of these data using PCA, and we observed that the PD assay using the CSDC2 and Resin only (PD-control) data clustered separately (Fig. S6). Next, we calculated a log_2_ ratio between PD and the input data set and, considered only positive interactions at the TOP 5% ranked genes for both PD-CSDC2 and PD-control. After that, we excluded the RNAs present in the TOP 5% of PD-control data to find the RNA present in cardiac mesoderm that interacts with CSDC2 (Fig. 5A-B).

**Fig. 5.**
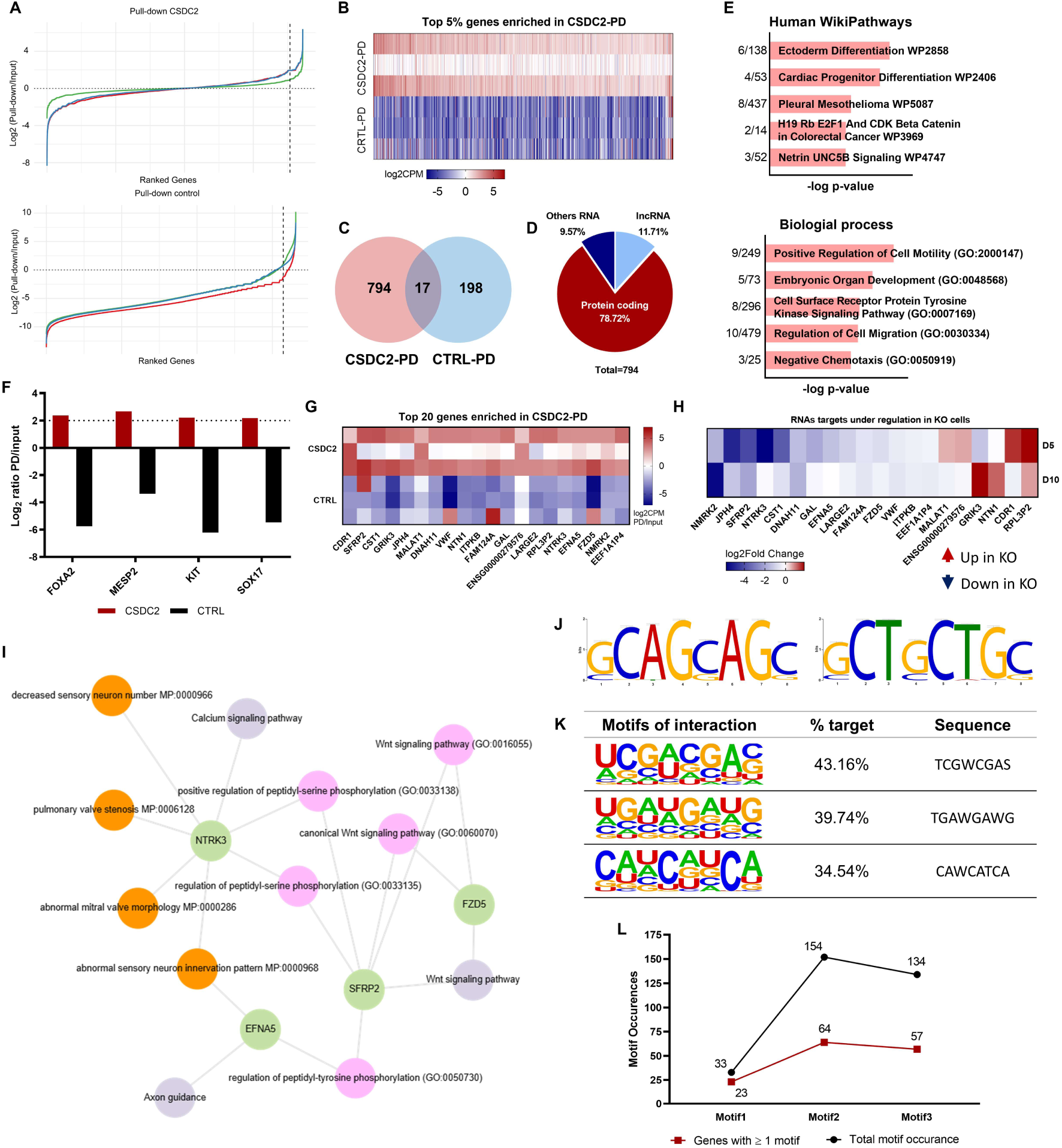
Description of CSDC2 RNA targets identified by pull-down assay. **A.** Enrichment analysis of genes in the CSDC2 pull-down (upper panel) and pull-down control (lower panel). The y-axis represents the log2 ratio relative to the input data. Dashed lines indicate the threshold of the top 5% most enriched genes in each analysis. **B.** Heatmap of the top 5% enriched genes in the CSDC2 pull-down dataset. Values are presented as the ratio of pull-down to input normalized read counts. The color key is displayed below the chart. **C.** Venn diagram of the top 5% enriched genes in the CSDC2 pull-down (PD) and control pull-down (CTRL-PD). **D.** Part-of-whole chart showing the biotypes of RNAs identified by CSDC2 pull-down. The percentage of each biotype is displayed next to the chart. **E.** Gene Ontology analysis of human WikiPathways (upper panel) and biological processes (lower panel) using RNAs identified by CSDC2 pull-down. **F.** Enrichment of cardiac progenitor genes in the CSDC2 pull-down and control pull-down datasets. The color key is shown below the chart. **G.** Heatmap of enrichment of the top 20 genes in the CSDC2 pull-down. Gene names are displayed below the chart. Values are presented as the ratio of pull-down to input normalized read counts. The color key is displayed on the right side of the chart. **H.** Heatmap showing enrichment of the top 20 CSDC2 targets in the polysomal fraction of hiPSC WT and CSDC2 KO at cardiac mesoderm (day 5) and cardiac progenitor (day 10). Values are presented as log2 fold change of hiPSC WT relative to CSDC2 KO. The color key is displayed below the chart. **I.** Regulatory network of CSDC2 targets. The network was constructed using EnrichR-KG with the top 20 identified targets. Gray circles represent the KEGG 2021 human database, pink circles represent GO Biological Process 2021, orange circles represent MGI Mammalian Phenotype Level 4 2021, and green circles represent the identified targets. **J.** Prediction of RNA motif for CSDC2 interaction using the MEME suite web tool. Normal sequence (left panel), reverse complement (right panel). **K.** Prediction of RNA motifs for CSDC2 interaction using the HOMER software. The percentage of targets and sequences is displayed in the table. **L.** Occurrence of predicted interaction motifs among the top 100 targets identified by CSDC2 pull-down. Quantification was performed using FIMO from the MEME suite. The red line represents genes containing at least one motif, and the black line represents the total number of motif occurrences.

We found 794 genes identified only in CSDC2 samples, and 78% of which are categorized as coding genes (Fig. 5C-D). GO analysis of these gene candidates confirmed the development relevance of CSDC2, showing terms like “Embryonic organ development”, “Positive regulation of cell motility” in the biological process, while the “Cardiac progenitor differentiation” appeared in the GO of human pathways (Fig. 5E). Among the RNAs that interact with CSDC2, we found MESP2, SOX17, KIT and FOXA2, all genes involved with cardiac mesoderm and progenitor commitment (Fig 5F). In Fig. 5G, we highlighted the top 20 genes found as enriched in CSDC2 samples. Then, we checked the dynamics of these 20 CSDC2 targets in our translatome data, and when comparing the WT and KO conditions, we found a down-regulated pattern of those genes (Fig. 5G-H). Because the identified targets were decreased in the CSDC2 KO translatome, these results suggest a possible mechanism of regulation by CSDC2 in the cardiac context. Next, we generated a network of molecular and biological processes using these genes and found that some targets act within the same functional group, with a strong enrichment of the WNT signaling pathway within the regulatory network (Fig. 5I).

Finally, we used the pulldown dataset to identify consensus sequences among the candidate targets and to describe a potential RNA motif for CSDC2 interaction. First, we used the Multiple EM for Motif Elicitation (MEME) algorithm to identify motifs enriched in the top 100 genes identified by the pull-down assay. Using this approach, we found one putative motif of interest for CSDC2 interaction (Fig. 5J). In addition, we searched for enriched motifs using the Hypergeometric Optimization of Motif EnRichment (HOMER) tool applied to peaks from PD-RNA-seq, which led to the identification of three putative motifs, each present in more than 34% of the dataset (Fig. 5K). Next, we applied an additional tool to verify the occurrence of these motifs within the top 100 most enriched genes from the pulldown assay. Although motif 1 (TCGWCGAS) showed a higher percentage of targets in the initial algorithm, it appeared to be less prevalent among the most highly enriched candidate targets (top 100). In contrast, motif 2 and motif 3 displayed a higher overall occurrence, both in terms of total occurrences and number of genes containing the motif (Fig. 5L). Together, these two motifs represent strong candidates for future studies on CSDC2–RNA interactions.

## Discussion

RNA-binding proteins (RBPs) are key regulators of post-transcriptional gene expression. They play a crucial role in human development, and any dysregulation can lead to pathological processes^38,39^. In the context of cardiovascular health, several RBPs have been linked to a range of cardiovascular diseases (CVD), such as cardiomyopathy, heart failure, and diabetic vascular disease^40–43^. These conditions are among the leading causes of death worldwide^8^. In recent years, researchers have focused on targeting these RBPs with specific molecules to develop new therapeutic strategies for RBP-related diseases^44,45^. These data highlight the need for characterizing RBPs involved in cardiovascular development and homeostasis. Here, we describe Cold Shock Domain Containing C2 (CSDC2) as an RBP that participates in cardiac differentiation using a human pluripotent stem cell (hPSC) model. We found that the absence of CSDC2 impairs cardiomyocyte differentiation in these cells, strongly suggesting an essential role in the differentiation process. Additionally, we have mapped its interactome and identified several RNAs that CSDC2 may interact with in the context of cardiac development and function.

CSDC2 (also called PIPPin) was initially described as an RBP expressed in the encephalon during rat development, enriched in the cerebral cortex and Purkinje cells from the cerebellum^25,27,46^. Our previous data suggest that C*SDC2* expression also increases during the cardiac differentiation of hESC; however, the role of *CSDC2* in a cardiac context remains poorly explored. In this work, we showed the robustness of the expression profile of CSDC2 during the cardiomyogenic development model. *CSDC2* expression increases at the final stages of cardiac differentiation in both the embryonic body formation protocol and in a monolayer protocol, independent of the cell type used. We also showed that CSDC2 is present in cardiomyocytes derived from hiPSC. Indeed, we could not detect expression of CSDC2 in non-cardiomyocytes cells (e.g. smooth muscle cells, endothelial cells and fibroblast) in culture. CSDC2 has been described to be mainly expressed in brain-related cells^46^. However, it has also been described as not tissue-specific ^28–31^. In our model, CSDC2 was localized predominantly in the cytoplasm rather than in the nucleus in hiPSC-derived cardiomyocytes. The cellular localization of CSDC2 is related to its protein function. In neuron cells, CSDC2 is found only in the nuclear protein fraction^27^. On the other hand, during the decidualization process, the protein is present in the cytoplasm of tissue areas under differentiation. When the CSDC2 is in the nucleus, these areas were found under a proliferative state^28^. The mislocalization of an RBP may result in a pathological process, as reported for RBM20 in the cardiac context, which occurs in several types of dilated cardiomyopathy. In this scenario, the mislocalization of RBM20 results in an aberrant splicing process affecting the homeostasis in heart cells^47,48^.

In our model, loss-of-function of CSDC2 directly impacted in cardiomyocytes differentiation. CSDC2 knockout impairs the cells from differentiating into cardiomyocytes without affecting the differentiation into other cell types. We showed that the lack of CSDC2 altered NKX2-5 and TBX5 expression, two master regulators of cardiomyogenic development in humans ^49–51^. In fact, these two genes are known as master regulators of cardiac development, since they act as transcriptional factors responsible for activating the expression of cardiomyocyte genes, such as MLC2, NPPA, and MYH6, essential genes for heart functionality^52–55^. Our translatome data set shows an overall reduction in cardiomyogenic genes in knockout cells, while the expression of endoderm and ectoderm-derived genes increases. Knockout or mutated models have been employed to verify the relevance of these RBPs in heart development, as shown for several proteins; in fact, some proteins that present an altered phenotype were correlated with pathological processes such as QKI, RBM20, RBM24, and RBPMS^19,20,56–59^.

Although the expression profile of CSDC2 suggests the role of this protein in the final stages of cardiac differentiation, our data showed that its absence during the transition from mesoderm cardiac cells (day 5) into cardiac progenitor (day 10) had the greatest impact. When we observed the lack of differentiation of cardiac progenitors into cardiomyocytes, we hypothesized that recovering the expression of CSDC2 at days 5 or 7 could rescue the differentiation. Interestingly, only starting CSDC2 expression on day 5 rescued the differentiation capacity. This showed that even when CSDC2 maximum expression was detected in immature cardiomyocyte cells (day 15), its ability to regulate cardiac commitment happens at early stages of differentiation. These observations are supported by our findings about the interactome of CSDC2. We were able to describe two independent interactome patterns related to distinct biological processes. When CSDC2 is present in cardiac mesoderm cells, the interactome network suggests a role as a post-transcriptional regulator. On the other hand, in cardiac progenitor cells, CSDC2 interacts with proteins involved in cell cycle regulation. These findings could categorize CSDC2 as a moonlight protein, which participates in two physiology processes depending on its intracellular environment^60^. Other proteins have already been described as acting as moonlight proteins during cardiac development, such as PKM2, a kinase with a non-enzymatic role that is able to regulate the expression of genes such as *Cyclin D1* and *c-Myc*, as well as showing an enzymatic role that increases flux through the pentose phosphate pathway, resulting in changes in redox metabolism^61^.

The characterization of an RBP role in development and/or pathological process includes the identification of potential targets for this protein. Here, we also showed the potential interaction of CSDC2 with mRNAs described to participate in cardiac differentiation commitment. We wanted to highlight the presence of MESP2, SOX17, and FOXA2 among the identified mRNAs. MESP2 is described as participating in cardiac mesoderm differentiation, and its dysregulation can result in different pathological processes^62–64^. SOX17 also participates in cardiac mesoderm commitment, as well as in the specification of endocardium development^65,66^. FOX2A is present in cardiac progenitor cells and gives rise to cardiac ventricle cells^67^. Altogether, this suggests that CSDC2 could interact with mRNAs coding for crucial transcriptional factors able to regulate heart development. Among the top 20 genes identified through pull-down assays, we found SFRP2, FZDR5, JPH4 (isotype of JPH2, mainly expressed in heart muscle), and GRIK3—genes already known to participate in heart development or cardiopathies and in the WNT signaling pathway^68–71^. We can hypothesize that CSDC2 could stabilize or direct these mRNAs to the translation machinery, since some of them were down-regulated in translatome data of KO cells, and we also observed the interaction with ribosomal proteins. Finally, using bioinformatic tools and in silico analyses, we identified three putative RNA motifs present in the RNA that showed CSDC2 interaction. Further studies will be required to experimentally characterize CSDC2–RNA interactions, which was beyond the scope of the present study.

Although there is no animal model described for studying the phenotype of CSDC2 loss during development, low expression levels of CSDC2 in humans have been associated with a higher prevalence of heart failure^72,73^. Our knockout model showed that loss of CSDC2 reduces cardiac progenitor generation, providing insight into the protein’s function in humans. Since a reduction in cardiac progenitor cells could lead to fewer cells for heart development, this may impact the thickness of the heart wall, one of the causes of heart failure. Moreover, an RBP able to regulate cardiomyogenic process impacting in NKX2-5 and TBX5 expression deserves further attention. This work takes a step toward describing CSDC2 as an important RNA-binding protein that participates in post-transcriptional regulation in the cardiac context.

## Acknowledgments

We would like to thank Dr. Susanne Kramer (University of Würzburg, Germany) for providing the TurboID sequence. We would like to thank all the staff of the Carlos Chagas Institute (FIOCRUZ-PR) for the laboratory and administrative support. We also would like to thanks Program for Technological Development in Tools for Health (RPT-FIOCRUZ) for using the Genomic, Proteomic, qPCR and Microscopy facilities at Instituto Carlos Chagas – Fiocruz/PR.

## Data availability

RNA-seq data are available at the National Center for Biotechnology Information (NCBI) Sequence Read Archive, accession number PRJNA1413977 upon publication of this article. The proteomics data of this study are not openly available, but are accessible from the corresponding author upon request.

