## Supplementary material for "CSDC2, an RBP essential to cardiomyocyte commitment during cardiac differentiation in hiPSC"

### Gomes-Júnior, R. et al., 2026 - Supplementary material

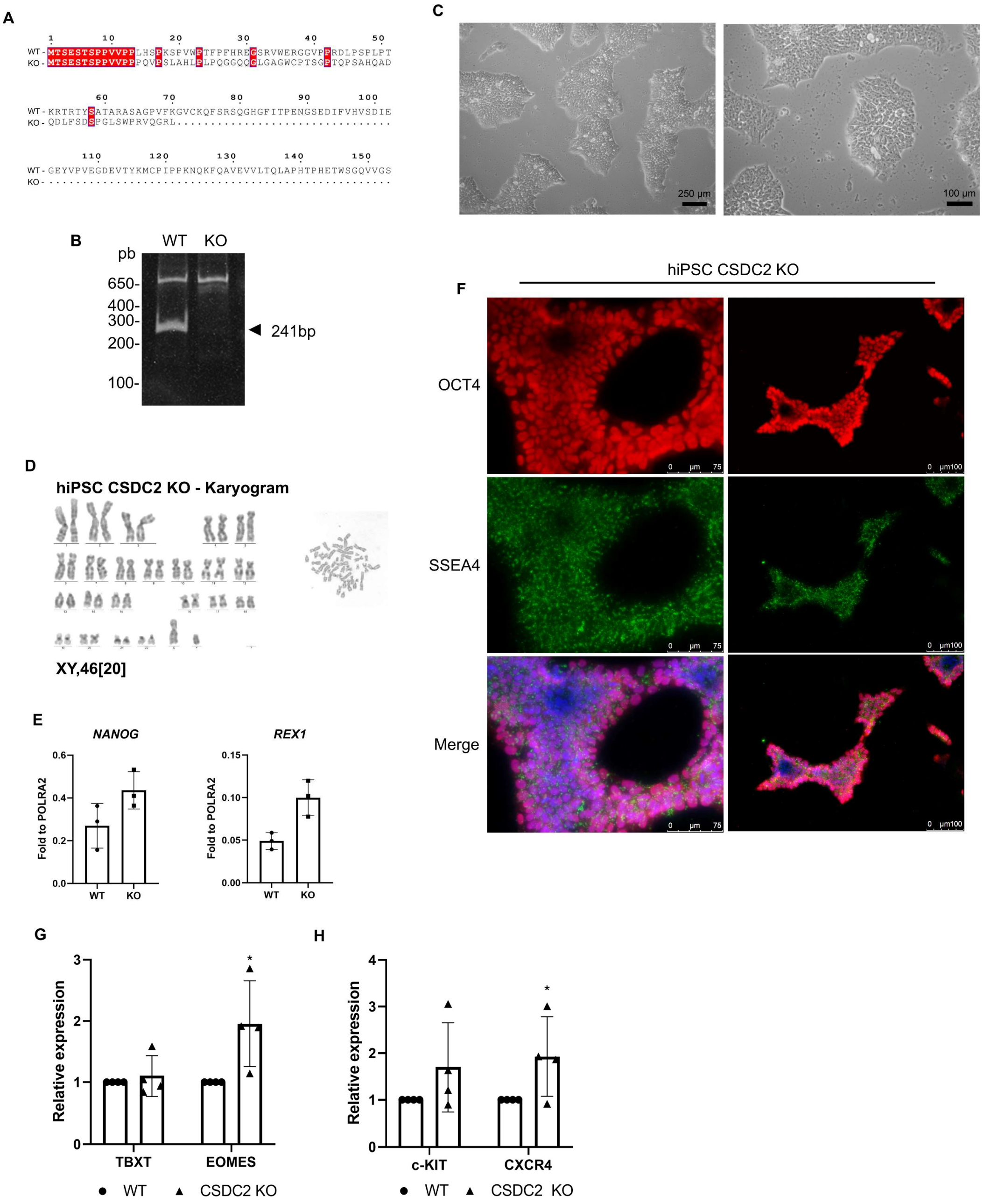


**Fig. S1. Characterization of the CSDC2 knockout hiPSC line. A.** Amino acid sequence alignment of the wild-type and CSDC2 knockout gene sequences. Identical residues are highlighted in red. **B.** Amplification of the CSDC2 genomic sequence using primers for the deleted region in hiPSC WT and CSDC2 KO cells by conventional PCR, separated on a 1% agarose gel. Arrowhead indicates the expected 241 bp band. **C.** Brightfield microscopy showing colony morphology of hiPSC CSDC2 KO cells. The scale bar is shown in the lower right corner. **D.** Karyogram of hiPSC CSDC2 KO. **E.** Relative expression of NANOG (left panel) and REX1 (right panel) in hiPSC WT and CSDC2 KO. **F.** Immunofluorescence of OCT4 and SSEA4 in hiPSC CSDC2 KO. OCT4 is shown in red, SSEA4 in green, and nuclei in blue. Images are shown at two magnifications: left panel (20×) and right panel (10×). **G.** Relative expression of mesoderm markers in hiPSC WT and CSDC2 KO at day 1 of the cardiac differentiation protocol, assessed by qPCR. **H.** Relative expression of cardiac mesoderm markers in hiPSC WT and CSDC2 KO at day 5 of the cardiac differentiation protocol, assessed by qPCR.

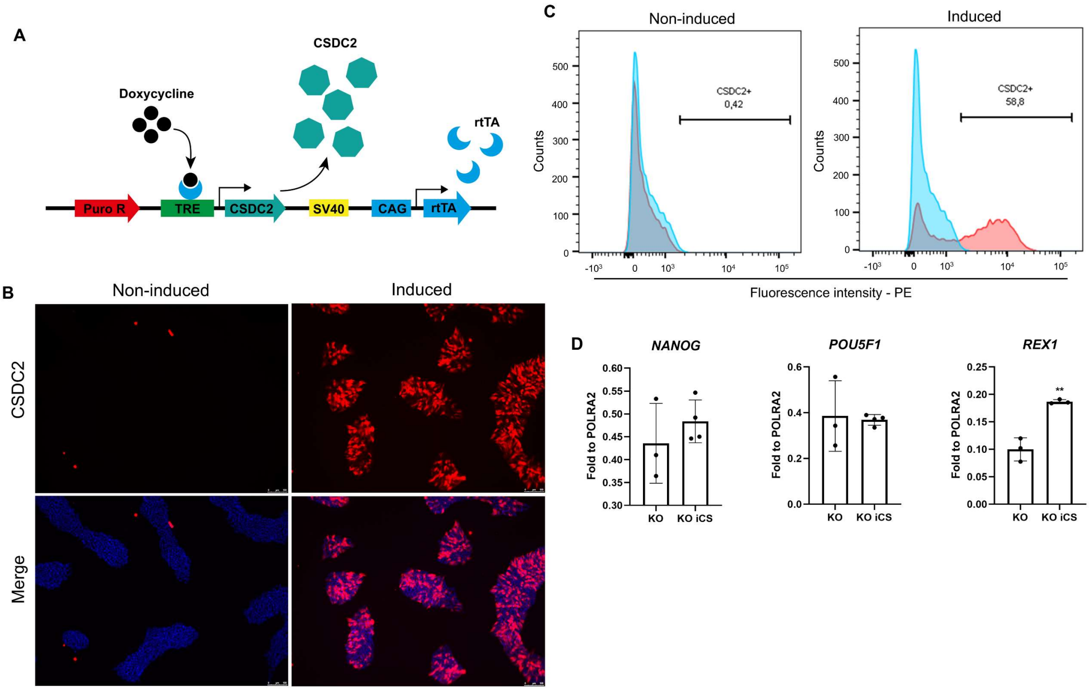


**Fig. S2. CSDC2-inducible expression cell line derived from knockout hiPSCs. A.** Scheme map of the CSDC2-inducible expression vector. **B.** Immunofluorescence of hiPSC KO iCSDC2. CSDC2 is shown in red and nuclei in blue. Non-induced (left) and induced (right). **C.** Flow cytometry analysis of cells responsive to the inducible system in hiPSC KO iCSDC2. Staining control is shown in blue, and stained cells are shown in red. Non-induced (left) and induced (right). The percentage of CSDC2-positive cells is indicated above the gate. **D.** Relative expression of pluripotency markers in CSDC2 KO and KO iCSDC2 in the pluripotent state, assessed by qPCR.

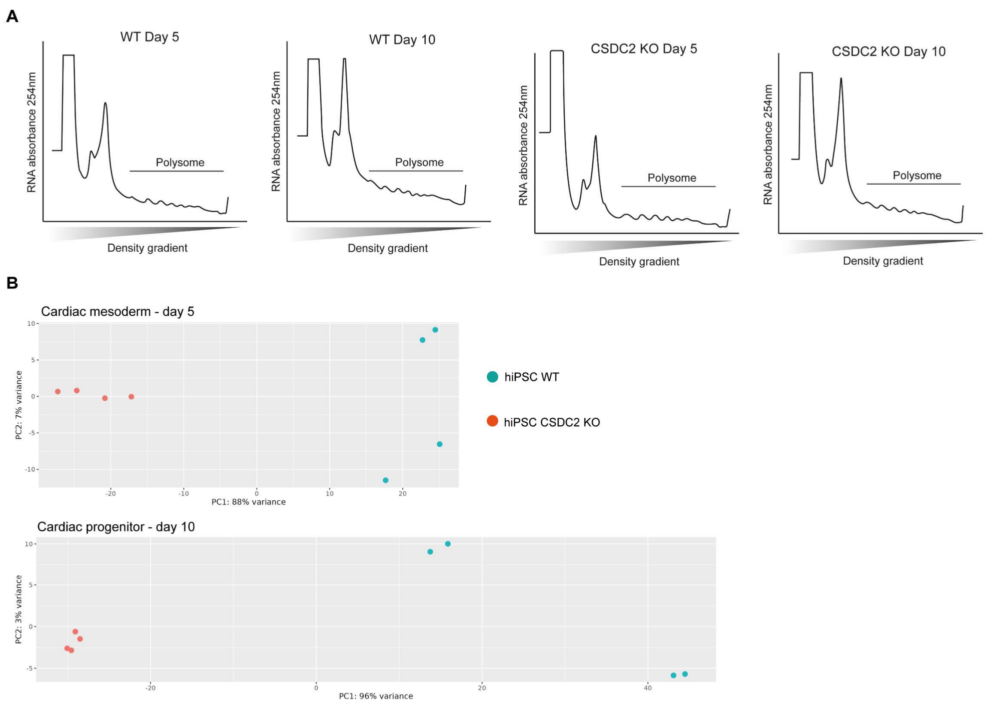


**Fig. S3. Polysome profile and translatome analysis. A.** Polysome profiles of hiPSC WT and CSDC2 KO at days 5 and 10 of cardiac differentiation. The y-axis represents RNA detection measured as absorbance at 254 nm, and the x-axis represents the density gradient. Polysome fractions are indicated in the chart. **B.** Principal component analysis (PCA) of RNA-seq data from the translatome dataset of cardiac mesoderm (upper) and cardiac progenitor (lower). The color legend is shown on right side of the chart.

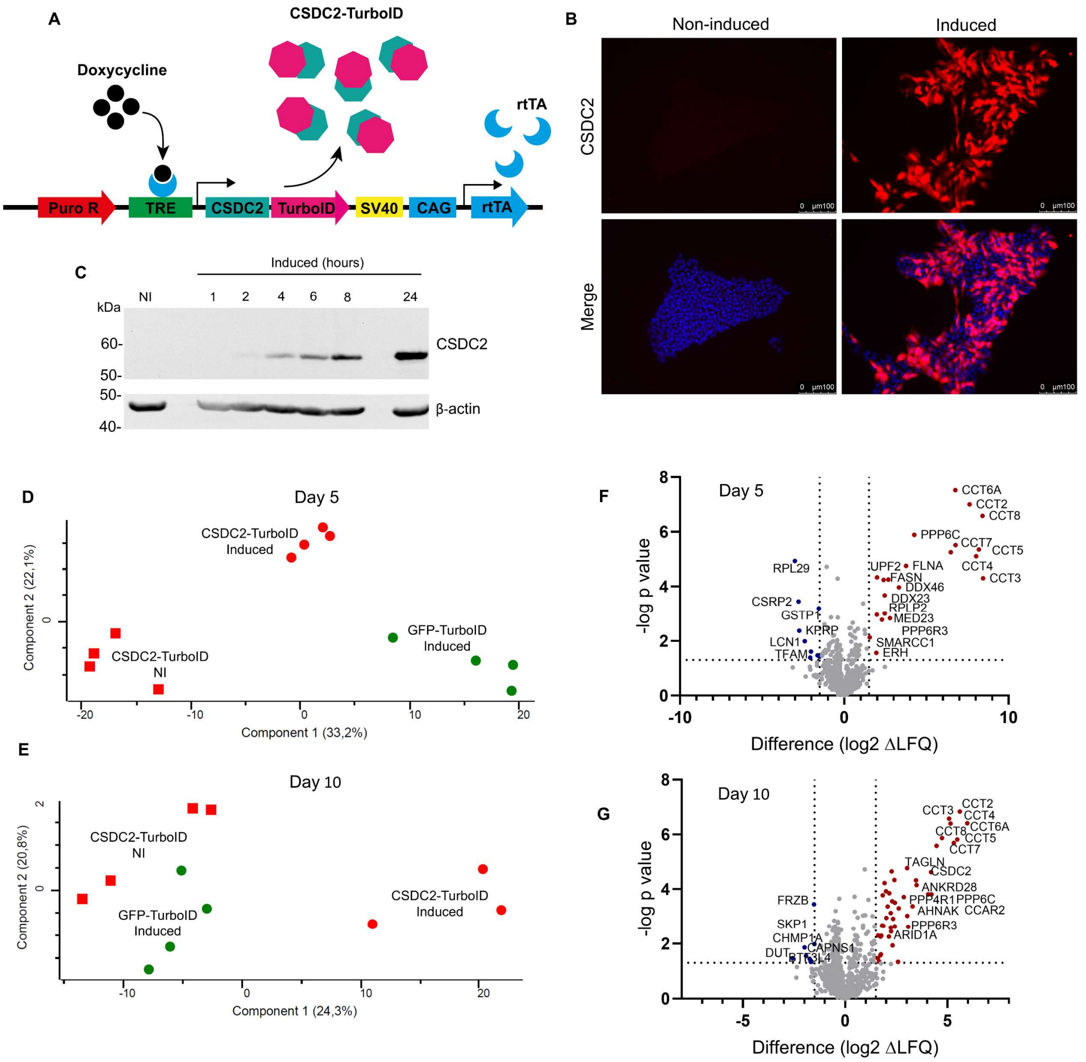


**Fig. S4. Generation of CSDC2-TurboID and proteomic data analysis of CSDC2 proximity labeling. A.** Scheme of the CSDC2-TurboID inducible system. **B.** Immunofluorescence of hiPSC iCSDC2-TurboID. CSDC2 is shown in red and nuclei in blue. Non-induced (left) and induced (right). **C.** Western blot analysis of hiPSC iCSDC2-TurboID at different induction times. The molecular weight ladder is shown on the right. Induction times are indicated above, and the antibodies used are indicated on the left. **D-E.** Principal component analysis (PCA) of the proteomic dataset in cardiac mesoderm (day 5) (D) and cardiac progenitor (day 10) (E) during cardiac differentiation using hiPSC iCSDC2-TurboID and hiPSC iGFP-TurboID. NI, non-induced; IND, induced. **F-G.** Volcano plots for cardiac mesoderm (day 5) (F) and cardiac progenitor (day 10) (G) showing data from induced and non-induced hiPSC iCSDC2-TurboID cells. Blue dots represent proteins enriched in the non-induced condition, while red dots indicate proteins enriched in the induced condition. Y-axis threshold: p-value ≤ 0.05 (−log p-value ≥ 1.3); x-axis threshold: log2 LFQ difference ≥ 1.5. Gene names are shown next to their representative dots.

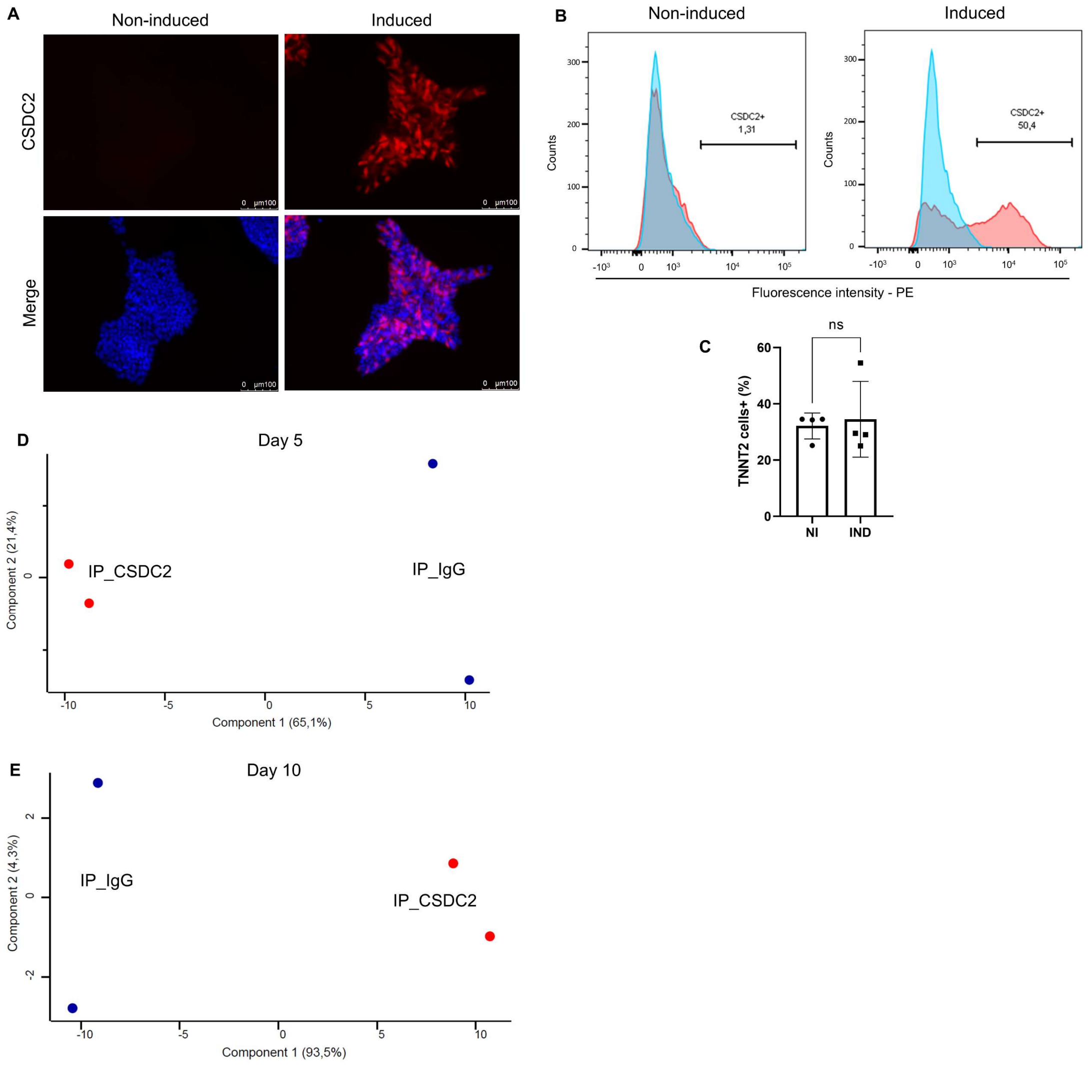


**Fig. S5. Generation of hiPSC CSDC2 overexpression and proteomic dataset analysis for CSDC2 immunoprecipitation. A.** Immunofluorescence of hiPSC iCSDC2. CSDC2 is shown in red and nuclei in blue. Non-induced (left) and induced (right). **B.** Flow cytometry analysis of cells responsive to the inducible system in hiPSC iCSDC2. Staining control is shown in blue, and stained cells are shown in red. Non-induced (left) and induced (right). The percentage of CSDC2-positive cells is indicated above the gate. **C.** Statistical analysis of cardiac differentiation efficiency in hiPSC iCSDC2. NI, non-induced; IND, induced. The y-axis represents the percentage of TNNT2-positive cells. **D–E.** Principal component analysis (PCA) of the immunoprecipitation proteomic dataset at the cardiac mesoderm (day 5) (D) and cardiac progenitor (day 10) (E) stages during cardiac differentiation. Experimental group names are shown next to the dots.

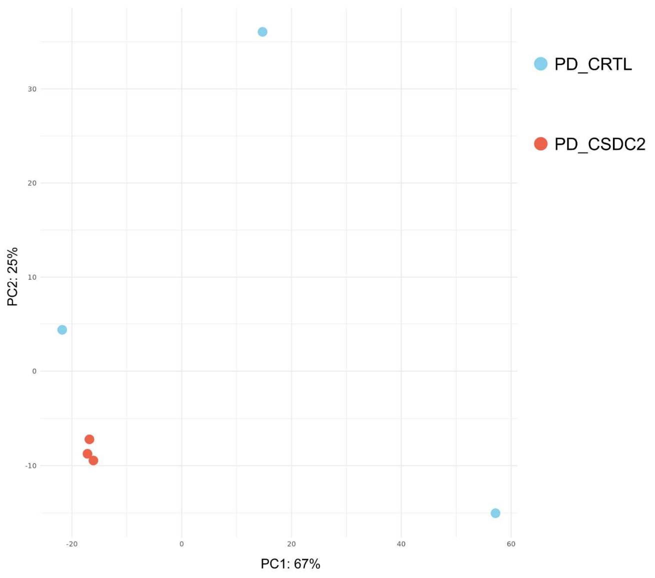


**Fig. S6. RNA-seq data analysis of CSDC2 pulldown.** Principal component analysis (PCA) of the pulldown RNA-seq data. The color legend is displayed on the right side.
